## Supplemental Data for "Structural basis for antibody cross-neutralization of Dengue and Zika viruses"

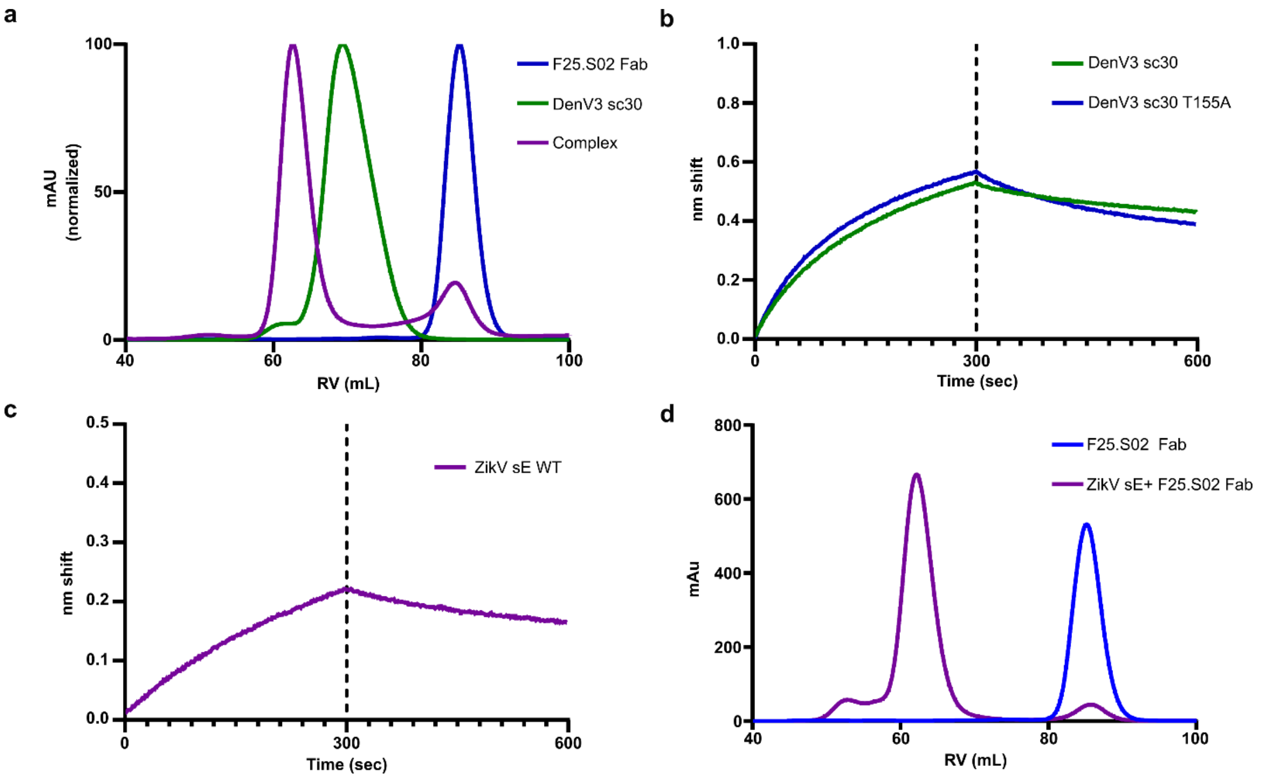


**Supplementary Figure 1**. **Purification and binding assays of F25.S02 Fab to DenV3 and ZikV sE dimers**. a) SEC traces of F25.S02 Fab, DenV3 sE sc30, and the complex of the two. mAu was normalized as F25 Fab expressed at a much higher level. b) BLI data of F25.S02 Fab binding to DenV3 sE sc30 and N153 glycan deletion. Binding data showed that presence of lack of glycan did not affect binding. c) BLI data of F25.S02 Fab binding to ZikV sE WT dimer. d) SEC traces of F25.S02 Fab and coexpression of ZikV sE WT + F25.S02 Fab

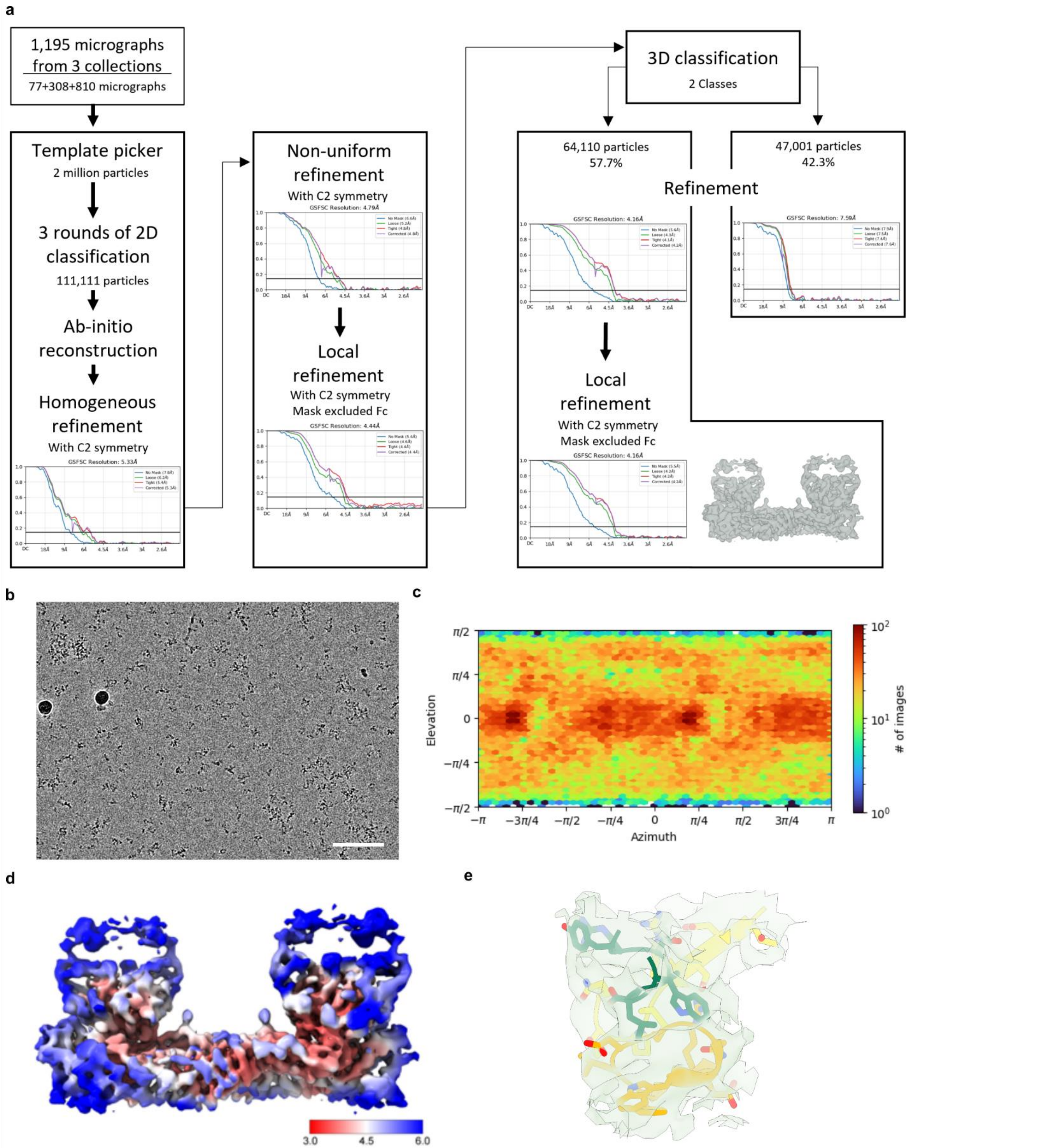


**Supplementary Figure 2**. **CryoEM data collection and processing**. a) Data processing workflow for DenV3 sE sc30 bound to F25.S02 structure. b) Representative motion corrected micrograph from data collection. Scale bar is 100 nm. c) View direction distribution plot of particles in 3D reconstruction. d) 3D reconstruction map colored by local resolution. e) Model fit image to 3D reconstruction. Focused on fusion loop and CDRH3 of F25.S02. Map is cut off within 3 Å of model.

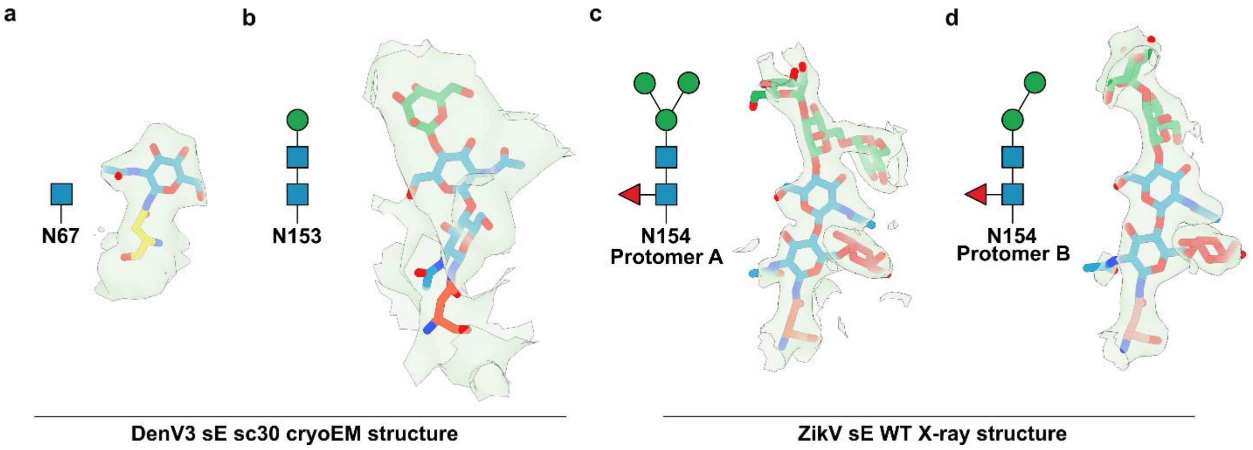


**Supplementary Figure 3. CryoEM map and electron density map around glycans.** a-b) Reconstructed cryoEM map within 3 Å of glycan at position 67 (A) and 153 (B) of DenV3. C2 symmetry was used in reconstruction so the map for glycans on each protomer is identical. c-d) Electron density contoured to 1.0 RMSD within 3 Å of the N154 glycan of ZikV. The glycan diagram for each is shown with blue squares, red triangles, and green circles representing N-acetylglucosamine, fucose, and mannose, respectively.

| | DenV3 sE sc30 bound to F25.S02 Fab |
|---|---|
| **Data collection** | |
| Microscope | Glacios |
| Voltage (kV) | 200 |
| Electron Dose (e-/Å$^2$) | 50 |
| Detector | K3 |
| Pixel Size (Å/px) | 1.122 |
| Defocus Range (µm) | -1.2 to -2.5 |
| Collection Tilt ($^o$) | 0 |
| Magnification | 36,000x |
| **Reconstruction** | |
| Software | CryoSPARC v4.4 |
| Selected Micrographs | 1,195 |
| Selected Particles | 64,110 |
| Symmetry | C2 |
| Box Size (px) | 320 |
| Resolution (Å) (FSC$_{0.143}$) | 4.16 |
| **Refinement** | |
| Map B factor (Å$^2$) | 215 |
| No. atoms* | 9,710 |
| Protein | 9,606 |
| Water | - |
| Ligand | 104 |
| Mean B-factor (Å) | |
| Protein | 137.3 |
| Water | - |
| Ligand | 130.4 |
| RMS bond length (Å) | 0.002 |
| RMS bond angle ($^o$) | 0.561 |
| **Validation** | |
| MolProbity[a] | 1.75 |
| Clashscore | 5.03 |
| CaBLAM outliers (%) | 3.75 |
| EMRinger[b] | 1.30 |
| Rotamer Outliers (%) | 0.95 |
| Ramachandran | |
| Favored (%) | 92.16 |
| Disallowed (%) | 0.00 |
| **PDB ID** | **9OWE** |
| **EMDB ID** | **70931** |

**Supplementary Table 1. CryoEM data collection and refinement statistics.** *Only none-hydrogen atoms. [a]Determined using MolProbity [1]. [b]Determined using EMRinger [2] in Phenix [3].

| | ZikV sE bound to F25.S02 Fab |
|---|---|
| **Data collection** | |
| Space group | $P4_12_12$ |
| Cell dimensions | |
| *a*, *b*, *c* (Å) | 117.86, 117.86, 346.10 |
| *α, β, γ* (°) | 90, 90, 90 |
| Resolution (Å) | 50.00-2.3 (2.43-2.30) |
| $R_{merge}$ [a] | 0.227 (1.089) |
| <I/σ(I)> | 15.19 (1.11) |
| $CC_{1/2}$ | 0.996 (0.482) |
| Completeness | 99.5 (99.8) |
| Redundancy | 7.66 (7.08) |
| **Refinement** | |
| Resolution (Å) | 48.70-2.30 (2.32-2.30) |
| No. unique reflections | 206,169 (33,462) |
| $R_{work}$ [b] / $R_{free}$ [c] | 0.225/0.274 (0.307/0.336) |
| No. atoms | 13,151 |
| Protein | 12,582 |
| Water | 1,006 |
| Ligand | 255 |
| B-factors (Å$^2$) | 68.64 |
| Protein | 68.83 |
| Water | 58.38 |
| Ligand | 84.20 |
| RMS bond length (Å) | 0.004 |
| RMS bond angle (°) | 0.75 |
| **Ramachadran Plot Statistics** [d] | |
| Residues | |
| Most Favored region | 96.65 |
| Allowed Region | 2.92 |
| Disallowed Region | 0.43 |
| Clashscore | 6.62 |
| **PDB ID** | **9OWF** |

**Supplementary Table 2. X-ray data collection and refinement statistics.** Numbers in parenthesis represent highest resolution shell. [a]$R_{merge} = [\sum_h \sum_i |I_h - I_{hi}| / \sum_h \sum_i I_{hi}]$ where $I_h$ is the mean of $I_{hi}$ observations of reflection *h*. [b] $R_{factor}$ and [c] $R_{free} = \sum ||F_{obs}| - |F_{calc}|| / \sum |F_{obs}|$ x 100 for 95% of recorded data ($R_{factor}$) or 5% data ($R_{free}$). [d] Determined using MolProbity [1].
